# Photosynthetic response of *Chara braunii* towards different bicarbonate concentrations

**DOI:** 10.1101/2023.02.08.527653

**Authors:** Carolin Magdalene Heise, Martin Hagemann, Hendrik Schubert

**Affiliations:** Institute of Biosciences, Department of Aquatic Ecology, University of Rostock, Rostock, Germany; Institute of Biosciences, Department of Plant Physiology, University of Rostock, Rostock, Germany

## Abstract

A variety of inorganic carbon acquisition modes have been proposed in Characean algae, however, the specific inorganic carbon uptake mechanism is unknown for the genus *Chara*. In the present study, we analyzed if *C. braunii* can efficiently use HCO_3_^-^ as a carbon source for photosynthesis. For this purpose, *C. braunii* was exposed to different concentrations of NaHCO_3_^-^ for different time scales. The photosynthetic electron transport through photosystem I (PSI) and II (PSII), the maximal electron transport rate (ETR_max_), the efficiency of the electron transport rate (α, the initial slope of the ETR), and the light saturation point of photosynthesis (*E_k_*) were evaluated. Additionally, pigment contents (chlorophyll *a*, chlorophyll *b*, and carotenoids) were determined. Bicarbonate addition positively affected ETR_max_ after direct HCO_3_^-^ application of both PSII and PSI, but this effect decreased after 1 h and 24 h. Similar trends were seen for *E*_k_, but no significant effect was observed for α. Pigment contents showed no significant changes in relation to different HCO_3_^-^ concentrations. To evaluate if cyclic electron flow around PSI was involved in active HCO_3_^-^ uptake, the ratio of PSI ETR_max_/PSII ETR_max_ was calculated but did not show a distinctive trend. These results suggest that *C. braunii* can utilize NaHCO_3_^-^ in short time periods as a carbon source but relies on other carbon acquisition mechanisms over prolonged time periods. These observations differentiate *C. braunii* from other examined *Chara* spp. and suggest a minor direct role of HCO_3_^-^ as a carbon source for photosynthesis in this alga.

## Introduction

At least 2.7 billion years ago, the biosphere became increasingly oxygenated during the so-called “Great Oxidation Event”. Central players in this transformation were cyanobacteria, the only prokaryotes performing oxygenic photosynthesis (Badger & Price, 2003; Shestakov & Karbysheva, 2017). Another 1.5 billion years later, eukaryotic green algae lead to the next massive changes in the Earth’s ecosystem (Raven et al., 2012; Bachy et al., 2022). Approximately 500 million years ago, embryophytes (more broadly known as land plants) evolved from a single streptophyte algal ancestor, once again leading to major global biogeochemical changes and ultimately to terrestrialization (Raven et al., 2012; Bowman, 2022; Fürst-Jansen et al., 2022). These changes in the biosphere greatly affected aquatic photosynthetic organisms and lead to specific adaptations, allowing these algae to survive in habitats with drastically declined inorganic carbon (Ci) availability.

Under current atmospheric conditions, aquatic systems are usually depleted in carbon dioxide (CO_2_) due to its substantially slower diffusion rate in water than in air. Furthermore, the solubility of CO_2_ depends on temperature and pH, which strongly influences the availability of different Ci species in the medium (Badger & Price, 2003; Beilby et al., 2022). At a pH of 8, the predominant Ci species is the poorly membrane-permeable bicarbonate (HCO_3_^-^), which is found to be a strong limiting factor for growth within most aquatic habitats (van den Berg et al., 2002; Schubert et al., 2017). Consequently, the possibility to use HCO_3_^-^ as a carbon source for photosynthesis is often discussed to be advantageous for aquatic organisms to circumvent the slow diffusion rates of CO_2_ (van den Berg et al., 2002; Badger & Price, 2003). Many photosynthetic organisms, including cyanobacteria and eukaryotic algae, evolved specialized CO_2_-concentrating mechanisms (CCMs) to overcome this limitation.

One well-described CCM exists in the unicellular freshwater algae *Chlamydomonas reinhardtii*. This green alga accumulates HCO_3_^-^ inside the cell and converts it into CO_2_ via carbonic anhydrases (CAs) in chloroplasts. By compartmentalizing the enzyme that catalyzes the process of CO_2_ fixation, ribulose 1,5-bisphosphate carboxylase/oxygenase (RubisCO) within a micro-compartment called pyrenoid and raising the local concentration of CO_2_ in its vicinity the efficiency of photosynthesis can be enhanced (Mackinder, 2018). An analogous compartment known as carboxysome has first been described in cyanobacteria in 1956 by Drews & Niklowitz. Further evidence for the occurrence of pyrenoid-like compartments in chloroplasts has been obtained in several groups of algae, including Streptophyta and subclades of the Phragmoplastophyta, the Charophytes (Graham et al., 1991; Domozych & Bagdan, 2022). And even though the existence of a pyrenoid is not a prerequisite for a CCM in eukaryotic algae, as shown by Goudet et al. (2020), it is discussed as a direct indication for the operation of a CCM. This mechanism permits many aquatic organisms to circumvent the present CO_2_ limitations because passive diffusion of CO_2_ is unable to meet the photosynthetic demands of complex eukaryotic photoautotrophs (Colman et al., 2002; van den Berg et al., 2002). Another way to drive Ci fluxes for photosynthesis inside algae are pH variations in the external medium, across membranes or compartments (Beilby & Bisson, 2012; Mackinder, 2018). The acidification of the medium or excretion of CAs can force the equilibrium state towards CO_2_ and thus facilitate passive diffusion (Beilby et al., 2022).

Many hypotheses regarding the Ci acquisition of *Charophyceae* include the active transport of HCO_3_^-^ across the plasma membrane. However, several authors have discussed the lack of direct demonstration (Beilby et al., 2022; Walker et al., 1980). This raises questions about the nature of a possible CCM in *Charophyceae*. Several Ci acquisition strategies have been proposed to increase CO_2_ concentrations inside the cell. They include: 1) cell-level proton pumps which are likely involved in the generation of acidic bands, thus facilitating passive diffusion of CO_2_ (Walker et al., 1980); 2) enzymes like CAs that convert HCO_3_^-^ into CO_2_ and cooperate with H^+^/HCO_3_^-^ co-transport (Price et al., 1985); 3) specialized cellular structures called charasomes involved in Ci acquisition (Domozych & Bagdan, 2022). It has been shown that charasomes are sides of particularly intensive ion fluxes (Beilby & Bisson, 2012; Franceschi & Lucas, 1980; Hoepflinger et al., 2017; Pertl-Obermeyer et al., 2018). Their formation is light-dependent and they are certainly involved in the stabilization of acidic zones surrounding *Chara* cells (Foissner et al., 2015; Absolonova et al., 2018; Pertl-Obermeyer et al., 2018). These findings support the hypothesis that charasomes might facilitate Ci uptake for photosynthesis (Eremin et al., 2019).

Recent studies showed that photosynthesis in *Chara* spp. is often accompanied by calcification, especially at high pH conditions. These authors investigated 7 corticated *Chara* species, *C. aculeolata, C. aspera, C. globularis, C. hispida, C. tomentosa, C. vulgaris*, and *C. subspinosa*, regarding photosynthesis and calcification (Sand-Jensen et al., 2018; Ray et al., 2003). Additional results are available for *C. corallina*, focusing on transmembrane ion currents and HCO_3_^-^/OH^-^ transport across its plasma membrane (Lucas, 1975; Lucas & Nuccitelli, 1980), and *C. glomerata*, in which the co-transport of HCO_3_^-^/H^+^ has been analyzed (Choo et al., 2002). Furthermore, alternating low and high pH bands are believed to support Ci uptake in *C. australis* (Pertl-Obermeyer et al., 2018). Collectively, many studies postulated the efficient use of HCO_3_^-^ for photosynthesis by several *Chara* species.

Due to the available genome sequences (Nishiyama et al., 2018), *Chara braunii* can be regarded as an important model organism among *Charophyceae*. Here we analyzed which Ci acquisition strategy allows *C. braunii* to grow photosynthetically in low Ci environments because no physiological investigations of the Ci acquisition mode exist for this emerging model organism. In the present study, the photosynthetic performance of *C. braunii* was monitored under different bicarbonate (HCO_3_^-^) levels by Dual-PAM measurements (PSI and PSII) that permitted transient *in vivo* analyses of its immediate effects on photosynthesis.

## Material and Methods

### Sampling and culture conditions of *Chara braunii*

The charophytes used in this study were obtained from natural sediment collected from Lausiger Teiche (Saxony-Anhalt, Germany) by H. Schubert in January 2022. This site is known to be prone in charophyte vegetation and the occurrence of *C. braunii* has previously been observed (Becker et al., 2016). Approximately 100 mL of natural sediment were transferred to 0.5 L beakers and covered with 400 mL filtered (55 μm gaze) habitat water. *C. braunii* grew and sprouted after approximately 6 weeks at 21°C with a light intensity of 50 - 60 μmol photons m^-2^ s^-1^ under a 16 h/8 h light/dark cycle. Upon sufficient growth, *C. braunii* was identified according to Becker et al. (2016).

### Short-term exposure treatment

The photosynthetic response of *C. braunii* to elevated bicarbonate levels was analysed by incubating the top three whorls of 5 ≥ separate algae in 50 mL solutions of increasing NaHCO_3_^-^ concentrations for three different time points. *C. braunii* was exposed to elevated HCO_3_^-^ levels for 1 hour (h) or 24 h. In addition, increased NaHCO_3_^-^ amounts were directly applied before the measurements of photosynthetic parameters (0 h). For the longer incubation periods, the algae were placed in 50 mL falcon tubes and returned to the cultivation conditions before the measurement started. The following bicarbonate concentrations were applied: 100, 200, 500, and 1000 μM. For this purpose, a stock solution of NaHCO_3_^-^ was prepared (pH 7.8). To permit air contact during the time course of the experiment, the stock solution was stored in a tightly sealed glass bottle filled with at least to ¾ of the total volume. Filtered habitat water (55 μm gaze) served as a reference for the elevated concentrations and is referred to as “habitat”.

### Photosynthetic analysis

Photosynthesis was measured at room temperature with the DUAL-PAM-100 (Walz GmbH, Germany), allowing the parallel recording of photosystem II (PSII) with a high-performance PAM chlorophyll fluorometer, and photosystem I (PSI), with a dual wavelength absorbance spectrometer. Prior to the measurements, *C. braunii* was dark-adapted for 30 min and subsequently, light curve measurements were performed. Apical thalli consisting of at least three whorls were transferred to a quartz cuvette containing the respective media and the apical whorl was the focus of the measurements. Each photosynthesis-irradiance (PI) curve, consisted of 16 subsequent irradiation steps (47, 56, 69, 93, 119, 148, 190, 244, 313, 398, 496, 610, 763, 935, 1151, and 1472 μmol photons m^-2^ s^-1^). The time set for each irradiance was 60 s. Photosynthetic parameters were calculated by the Dual PAM software (Version 3.18, Heinz Walz GmbH, Germany). The maximal electron transport rate (ETR_max_), the initial slope (α), and the light saturation (*E_k_*) were calculated by fitting the PI curve to the model of Eilers and Peters (1988) with the least squares method by the Solver add-in in Excel (Microsoft Office, 2013). Subsequently, the pigment contents of the same samples were analysed.

### Pigment analysis

Pigment analyses were performed after photosynthesis measurements using the first to third whorl including internodes of *C. braunii* (n ≥ 5). Chlorophyll *a* (chl *a*), chlorophyll *b* (chl *b*), and carotenoids were extracted in 3 mL N, N-dimethylformamide for two weeks in darkness at - 20°C. The supernatant was carefully transferred into a 1 mL quartz cuvette without algae residues and the absorbance was measured at 480, 630, 647, 664, and 750 nm using a spectrophotometer (Perkin Elmer UV/VIS spectrometer Lambda 2). The pigment contents were calculated according to the equation given by Wellburn, 1994.

### Data analysis

All statistical analyses were performed with R (R Core Team 2022, version 4.2.2). For the statistical analyses the R packages readxl (Wickham & Bryan, 2022), nortest (Gross & Ligges, 2015), carData (Fox et al., 2022), car (Fox & Weisberg, 2019), FSA (Ogle et al., 2022) and dplyr (Wickham et al., 2022) were used. For tests of normality, Shapiro-Wilk tests were performed followed by Levene’s test for homogeneity of variance. Normal and homogeneous distributed data were analysed by ANOVA and significant differences between the groups were analysed by Tukey multiple comparisons of means with a 95% family-wise confidence level. In the case of non-normally distributed data, analyses were performed by Kruskal-Wallis rank sum test followed by the Conover test as a non-parametric post hoc test with adjusted *p*-values by the Bonferroni method. Significance levels were set to *p* < 0.05. For data visualization, the R packages ggplot2 (Wickham, 2016) and patchwork (Pedersen, 2022) were used, for the depiction of the compact letter display the package multcompView (Graves et al., 2019) was used.

## Results

To evaluate whether *C. braunii* can efficiently use HCO_3_^-^ as a carbon source for photosynthesis, we exposed the alga for different time scales to increasing bicarbonate concentrations. The impact of different NaHCO_3_^-^ amounts on PSI and PSII performances was measured using the DualPAM. From these measurements, the ETR_max_ (maximal electron transport rate), α (efficiency of the electron transport rate), and *E_k_* (light saturation point of photosynthesis) were calculated. Figure 1 summarizes these photosynthetic parameters of PSII after the addition of 100, 200, 500, or 1000 μM NaHCO_3_^-^. The experiment includes a measurement of these parameters after the same treatment with habitat water in which *C. braunii* grew prior to the experiments, as a reference.

**Figure 1:**
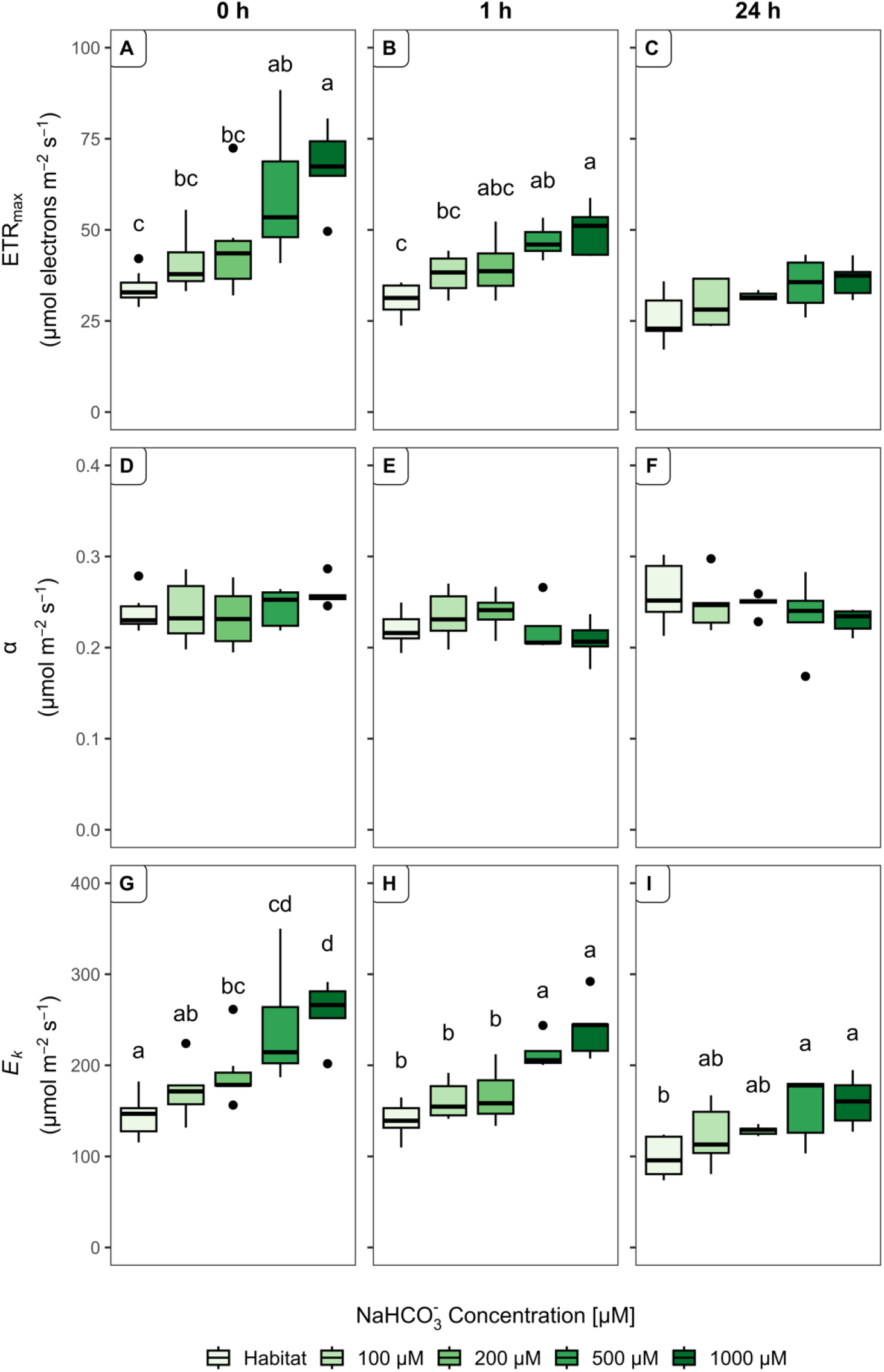
Bicarbonate effects on the PSII in *C. braunii*. Box plots of the effect of different NaHCO_3_^-^ concentrations on photosynthetic parameters of PSII including the maximal electron transport rate (ETR_max_), the initial slope of the photosynthesis-irradiance curve (*α*), and the saturation irradiance point for photosynthesis (*E*_k_). Values are shown for each evaluated time point after NaHCO_3_^-^ addition including the immediate effect (0 h), the effect after 1 hour (1 h), and after 24 hours (24 h). Box plots of n ≥ 5 stretch from the 25^th^ – 75^th^ percentile, black lines inside the boxes indicate the data median, and whiskers represent 1.5x the interquartile range limits. Outliers are shown as black dots. Equal letters connect homogeneous mean groups within the timepoint (Tukey HSD post hoc test, *p* < 0.05, Conover post hoc test, *p* < 0.05).

After direct application (0 h) of NaHCO_3_^-^ to the algae, a significant increase in PSII ETR_max_ with increasing concentration levels of NaHCO_3_^-^ was measured (ANOVA, *p* < 0.001). The most prominent difference was observed between the PSII ETR_max_ of *C. braunii* in habitat water conditions (no bicarbonate addition) compared to 500 and 1000 μM NaHCO_3_^-^ (Fig. 1A, Tukey’s test, *p* < 0.005). An almost three-fold higher PSII ETR_max_ was detected when the highest amount of bicarbonate was added prior to the measurement. After 1 h incubation at different bicarbonate levels, the positive effect of NaHCO_3_^-^ on PSII activity is still visible (Fig. 1B). However, the PSII ETR_max_ was generally less stimulated compared to the results obtained after the immediate application of the different bicarbonate concentrations. For example, supplementation with 1000 μM NaHCO_3_^-^ resulted in ETR_max_ close to 75 μmol electrons m^-2^ s^-1^ after direct bicarbonate addition, whereas ETR_max_ decreased to approximately 50 μmol electrons m^-2^ s^-1^ after 1 hour and even further to 30 - 40 μmol electrons m^-2^ s^-1^ after 24 h incubation time. In comparison to the reference measurements with habitat water, less varying electron transport rates at the three different time points were observed. i.e., ETR_max_ was 35 μmol electrons m^-2^ s^-1^ at 0 h and remained at 30 μmol electrons m^-2^ s^-1^ after 24 h. Thus, the changes in electron transport of PSII are HCO_3_^-^-dependent.

After 24 h incubation in elevated NaHCO_3_^-^ concentrations, the PSII ETR_max_ did not show any statistically significant differences between the bicarbonate supplemented groups including the habitat water reference (ANOVA, *p* > 0.05). Despite the general positive HCO_3_^-^-dependent effect on ETR_max_ values of PSII, this effect does not last over the time course of 24 h. NaHCO_3_^-^ supply did not appear to affect the efficiency of the electron transport rate (α) as this appears to be mostly unchanged among all measurements with the different bicarbonate concentrations at all time points (Figure 1 D-F). And even though the efficiency of the electron transport rate was not found to be NaHCO_3_^-^-dependent, the light demand for the saturation of PSII ETR_max_ increased with increasing NaHCO_3_^-^ concentrations, which could be taken as indication for a short-term active uptake of bicarbonate (*E*_k_, Figure 1G-I). After the immediate addition of NaHCO_3_^-^ (0 h) the increase of *E*_k_ is most prominent, showing a distinctive difference between the habitat water reference, 100 μM NaHCO_3_^-^ and 200 μM NaHCO_3_^-^ in comparison to 1000 μM NaHCO_3_^-^ (Figure 1G). A similar trend is still visible after 1 h supplementation with increasing NaHCO_3_^-^ concentrations (Figure 1H), clearly separating the light demand for saturation of the higher NaHCO_3_^-^ concentrations (500 μM and 1000 μM) from the lower concentration levels (100 μM, 200 μM, including the habitat water reference, ANOVA, *p* < 0.001). In contrast to PSII ETR_max_, the *E*_k_ increase by bicarbonate was still present in a significant manner after 24 h (Figure 1I, ANOVA, *p* < 0.05).

A similar experiment was conducted to analyse the response of photosynthetic parameters of PSI after the addition of different concentrations of NaHCO_3_^-^ (Fig. 2). Comparable differences for PSI as observed for PSII ETR_max_ can be seen for the immediate effects of NaHCO_3_^-^ (Fig. 2A), because a general increasing trend with increasing NaHCO_3_^-^ concentration was detected for PSI ETR_max_. The highest ETR_max_ values of PSI were observed at 1000 μM NaHCO3 ^-^ as observed for PSII. In the absorption measurements of PSI, less distinctive patterns are visible as previously observed for PSII, which is most striking 1 h after the addition of NaHCO_3_^-^ for the values of PSI ETR_max_ and *E*_k_ (Fig. 2B and H, compare with Fig. 1B and H). However, as for PSII no significant differences were observed among the α values of PSI at the different bicarbonate concentrations after the different incubation times (Fig. 2D-F, ANOVA/Kruskal *p* > 0.05).

**Figure 2:**
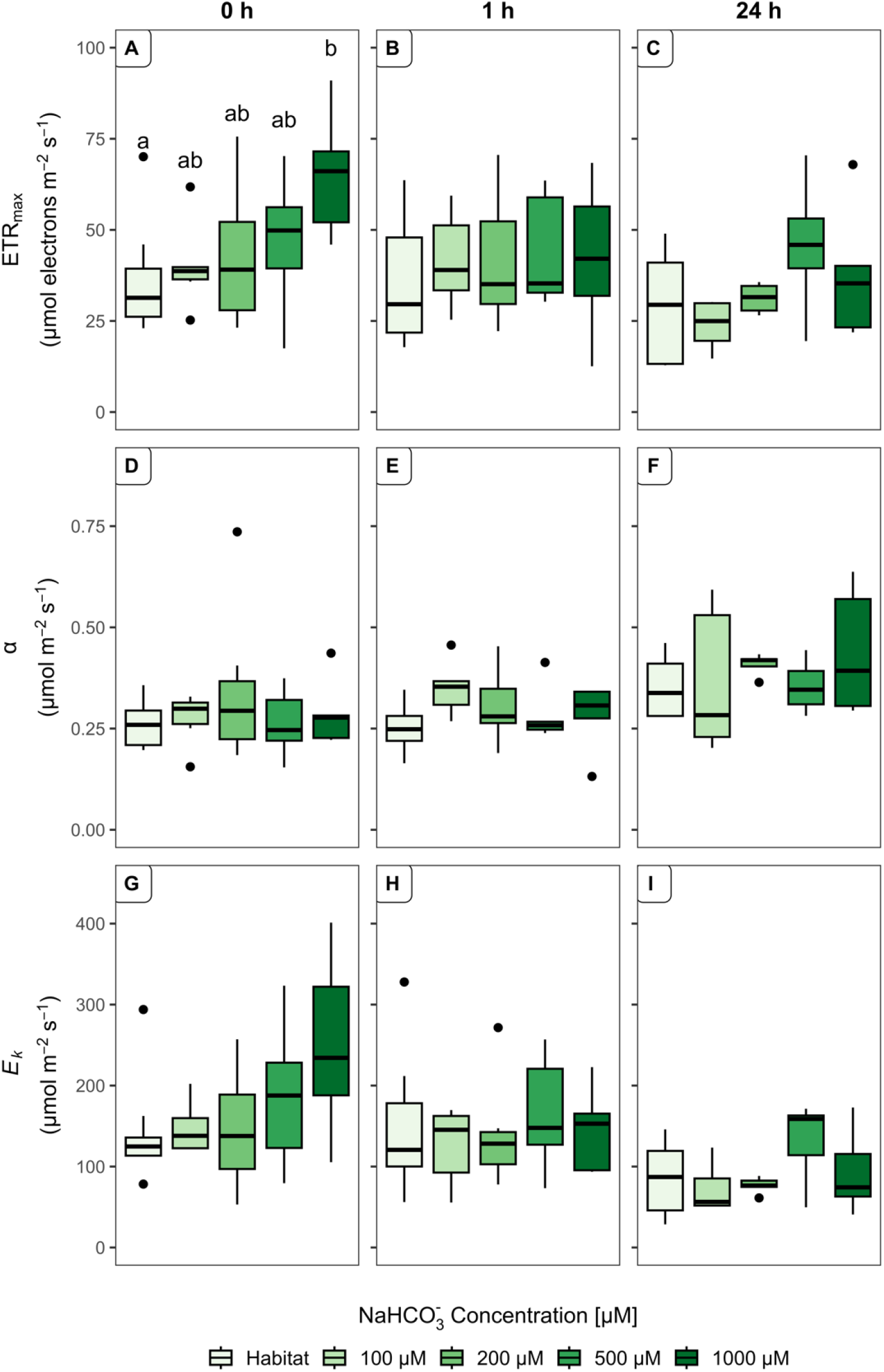
Bicarbonate effects on the PSI in *C. braunii*. Box plots of the effect of different NaHCO_3_^-^ concentrations on photosynthetic parameters of PSI including the maximal electron transport rate (ETR_max_), the initial slope of the photosynthesis-irradiance curve (*α*), and the saturation irradiance point for photosynthesis (*E*_k_). Values are shown for each evaluated time point after NaHCO_3_^-^ addition including the immediate effect (0 h), the effect after 1 hour (1 h), and after 24 hours (24 h). Box plots of n ≥ 5 stretch from the 25^th^ – 75^th^ percentile, black lines inside the boxes indicate the data median, and whiskers represent 1.5x the interquartile range limits. Outliers are shown as black dots. Equal letters connect homogeneous mean groups within the timepoint (Tukey HSD post hoc test, *p* < 0.05, Conover post hoc test, *p* < 0.05).

Analysis of *E*_k_, revealed increasing tendencies for light demand for the saturation of PSI ETR_max_, however, no statistically significant differences were observed. Furthermore, a similar trend in decreasing maximal values for PSI over bicarbonate exposure time can also be observed here, especially for *E*_k_. For example, the addition of 1000 μM NaHCO_3_^-^ resulted in maximal *E*_k_ values of around 380 μmol m^-2^ s^-1^ at time point 0 h, whereas the *E*_k_ values of PSI decreased to approximately 250 μmol m^-2^ s^-1^ after 1 h and around 170 μmol m^-2^ s^-1^ after 24 h, respectively.

Collectively, these results suggest that the photosynthetic efficiency of neither PSII nor PSI was influenced by different NaHCO_3_^-^ concentrations, however, clear immediate NaHCO_3_^-^- dependent stimulatory effects were observed for both ETR_max_ and *E*_k_ (Figs. 1 and 2). These effects decrease after 1 h and nearly vanished after 24 h among both parameters. These findings suggest that higher NaHCO_3_^-^ concentrations shortly raise the light demand for the saturation of ETR_max_ as reflected by significantly increased values of *E*_k_ from 100 μM to 1000 μM, but long-term incubation at elevated NaHCO_3_^-^ concentrations had no advantage for the photosynthetic activity of *C. braunii*.

Furthermore, the ratios of PSI ETR_max_/ PSII ETR_max_ were compared to assess the energy supply of *C. braunii* at increasing NaHCO_3_^-^ concentrations. In plants, an alternative electron flow is needed to increase the ATP/NADPH production ratio and satisfy energy supplies, which is achieved by the combined action of the linear electron flow (LEF) from PSII to PSI and the cyclic electron flow (CEF) around PSI (Wang 2022). Hence, we assumed that if *C. braunii* utilizes HCO_3_^-^ as a carbon source for photosynthesis an active uptake of the negatively charged ion is needed and this would be reflected in the ratio of PSI ETR_max_/ PSII ETR_max_. The respective calculation is depicted in Figure 3 for all three incubation times with NaHCO_3_^-^. Notably, the measurements of the habitat water reference were omitted from this analysis as the HCO_3_^-^ concentration here is unknown and cannot be properly arranged within the NaHCO_3_^-^ concentrations to visualize a trend in PSI ETR_max_/ PSII ETR_max_.

**Figure 3:**
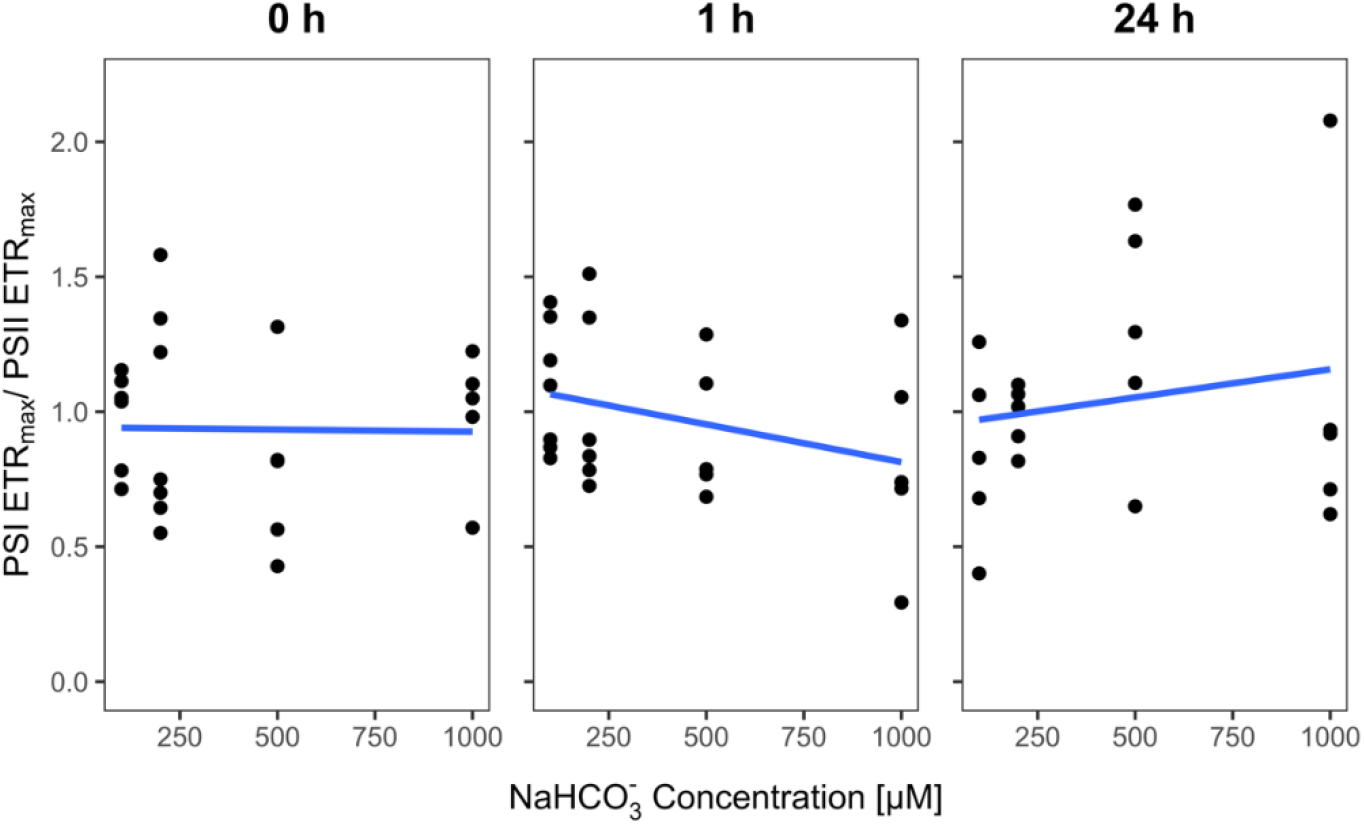
Bicarbonate effects on the cyclic electron flow in *C. braunii*. Estimated trend of cyclic electron flow (CEF) in *C. braunii* based on the maximal electron transport rate (ETR_max_) of PSI and PSII for each time point (direct addition (0 h), incubation for 1 h and 24 h) in NaHCO_3^-^_ concentrations of 100 μM, 200 μM, 500 μM, and 1000 μM. Black dots indicate separate measurements n ≥ 5. Trends are visualized by a regression line shown in blue.

After direct application of NaHCO_3_^-^ no change in PSI ETR_max_/ PSII ETR_max_ was observed (Fig. 3). After 1 h the ratio appears to decrease with increasing NaHCO_3_^-^ concentrations, while after 24 h an opposite trend can be observed. Even though it appears that the ratio PSI ETR_max_/ PSII ETR_max_ increases after 24 h it must be noted that this is based primarily because of one measurement in the 1000 μM NaHCO_3_^-^ concentration and, hence, this trend must be taken with caution. Overall, the PSI ETR_max_/ PSII ETR_max_ values appear to broadly differ in a range from 0.5 to 1.5 in nearly all cases. Compared to the obtained single parameters of PSI (Fig. 2) and PSII (Fig. 1) no reliable increase of PSI ETR_max_/ PSII ETR_max_ can be observed because in most cases just one measurement is responsible for the depicted trend and can be neglected. Thus, *C. braunii* does not seem to rely on CEF to fuel the uptake of HCO_3_^-^ across its membrane.

After the photosynthesis-irradiance bicarbonate curves were measured, pigment contents in *C. braunii* thalli were analyzed after the short-term exposure to NaHCO_3_^-^. The main photosynthetic pigments chlorophyll *a* (chl *a*), chlorophyll *b* (chl *b*), and carotenoids (car) were quantified. Commonly, the chl *a*/ chl *b* ratio is taken as an indicator of the photosynthetic properties and physiological status of the plant and is broadly used as an indicator of acclimation mechanisms (Küster et al., 2004; Gómez & Huovinen, 2011), while carotenoids were proposed to be photo-protective pigments in *Charophyceae* (Schagerl & Pichler, 2000). Hence, here the ratios of chl *a*/ chl *b* and chl *a*/ car were analyzed and are shown in Figure 4. No significant differences were observed for either chl *a*/ chl *b* or for chl *a*/ car. Steady pigment content trends can be observed across all NaHCO_3_^-^ concentrations, including the habitat water reference, and no significant changes are shown in proceeding time from direct application to 24 h.

**Figure 4:**
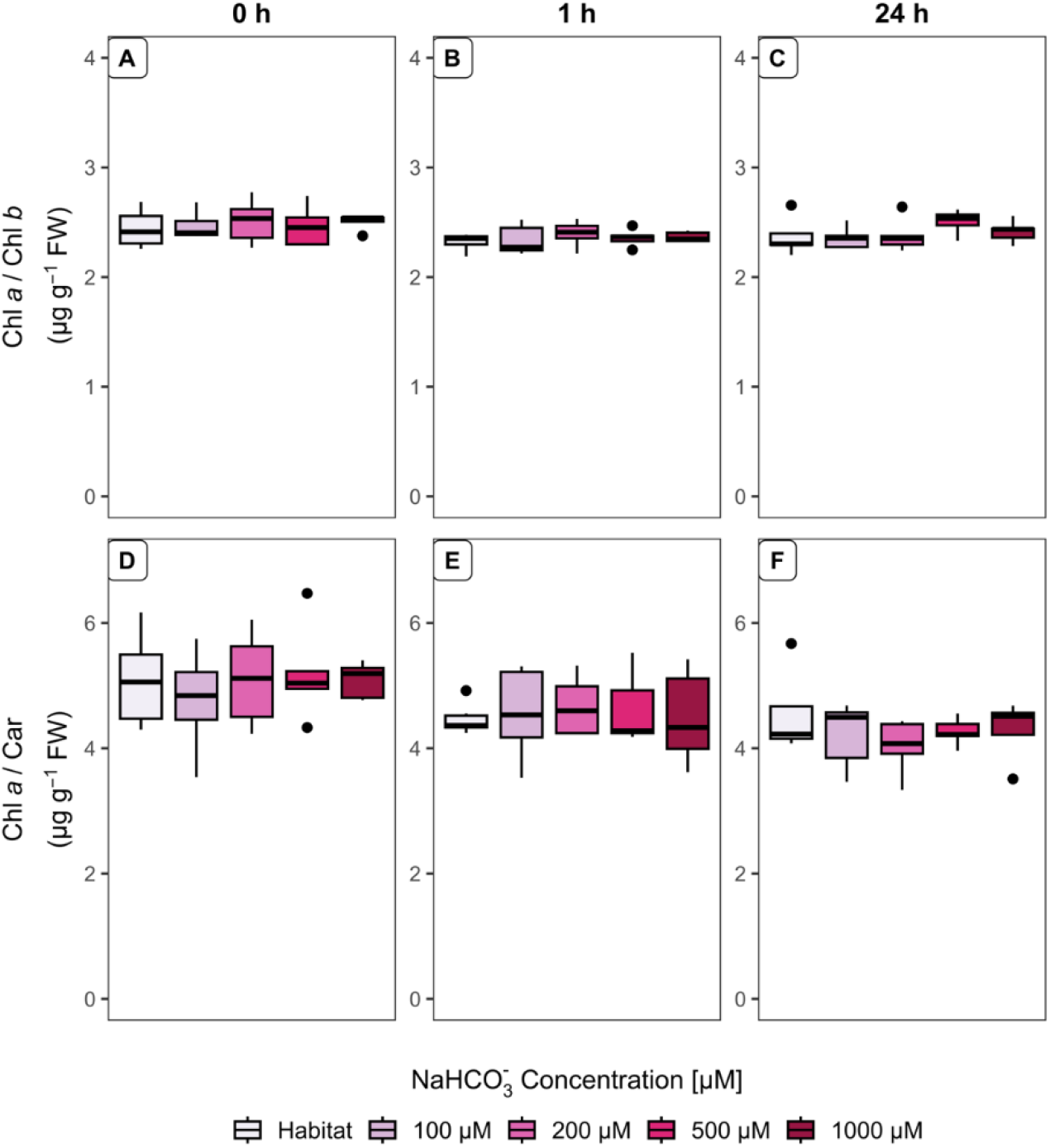
Bicarbonate effects on the pigmentation in *C. braunii*. Ratios of Chlorophyll *a* (chl *a*)/ Chlorophyll *b* (chl *b*) and chl *a*/ Carotenoid (Car) content after treatment of *C. braunii* with different NaHCO_3_^-^ concentrations and without treatment (Habitat). Box plots of n ≥ 5 stretch from the 25^th^ – 75^th^ percentile, black lines inside the boxes indicate the data median, and whiskers represent 1.5x the interquartile range limits. Outliers are shown as black dots.

## Discussion

In the last years, research focusing on Ci acquisition in non-model aquatic organisms steadily increased (Capó-Bauçà et al., 2022; Fürst-Jansen et al., 2022; Goudet et al., 2020; Magwell et al., 2023; Sand-Jensen et al., 2018; Zhou et al., 2016). These studies showed that further insights into the development of Ci acquisition mechanisms can only be obtained by analyzing species inhabiting different aquatic environments to assess evolutionary variations and gain a broader understanding of these adaptations to declining carbon environments (Capó-Bauçà et al., 2022).

The here presented results stand in contradiction to the previously widely accepted efficient use of bicarbonate by *Chara* spp. (Ray et al., 2003; Sand-Jensen et al., 2018), since our results reveal that *C. braunii* did not show long-lasting efficient use of NaHCO_3_^-^ as evaluated by photosynthetic parameters over the time course of 24 h. The short-term increase in ETR_max_ of PSII and PSI with increasing HCO_3_^-^ concentration levels, already decreased after 1 h, and no significant effects of HCO_3_^-^ were observed after 24 h (Fig. 1 and 2). These results suggest that a short-term uptake of bicarbonate can significantly increase the electron transport rate of photosynthesis, however, this effect does not withstand proceeding time periods, already starting at 1 h exposure. The presence of an ATP-dependent HCO_3_^-^ acquisition mechanism has previously been reported for *C. corallina* by Lucas et al. (1983). If a similar acquisition mechanism was present in *C. braunii*, the use of HCO_3_^-^ would require an active uptake mechanism to overcome the negative membrane potential (Beilby et al., 2022). One way to gain additional energy through ATP synthesis is the CEF around PSI (Wang et al., 2022). In this way, electrons are recycled and generate proton motive force without reducing NADP^+^ (Wang et al., 2022). This would consequently be reflected by the ETR ratio of PSI/PSII, with increasing values upon CEF presence (Jayasankar & Valsala, 2008). Our findings do not show an increase in PSI/PSII ratio of ETR with increasing NaHCO_3_^-^ concentrations and thus do not reflect an increase in energy supply by CEF (Fig. 3). However, the light demand *E*_k_ for the saturation of PSII ETR_max_ increased with increasing NaHCO_3_^-^ concentrations, which might indicate that products from the LEF are possibly used to fuel the bicarbonate uptake.

In previous studies, an immediate response of pigment contents was observed for *C. braunii* upon exposure to zinc and hydrogen peroxide (Herbst et al., 2021). These results suggest that a negative impact of elevated NaHCO_3_^-^ on the physiological status and pigmentation of the algae would already be prominent in short-term exposure experiments. Interestingly, Zhou et al. (2016) showed that elevated NaHCO_3_^-^ concentration has a negative effect on photosynthetic rate and pigment synthesis in macroalgae after 14 days. Such inhibitory effects on photosynthesis were not observed in the here presented study, and NaHCO_3_^-^ was not found to negatively affect pigment contents in *C. braunii*. Remarkably, the pigment content did only marginally change for chl *a*, chl *b*, and carotenoids with increasing NaHCO_3_^-^ concentrations.

Furthermore, no significant differences were observed in either of the applied exposure times at different NaHCO_3_^-^ concentrations (Fig. 4). The ratio between chl *a*/ chl *b* did not change, indicating a steady physiological status of *C. braunii* throughout the time periods of the experiment (Jayasankar & Valsala, 2008). The absence of an increase in carotenoids also points at a mild stress situation, because this pigment has been reported to play an important role in photo-protection in *Chara* (Küster et al., 2004).

Taken together our results suggest that NaHCO_3_^-^ can shortly be utilized by *C. braunii* however not for prolonged time periods and the nature of the energy supply is independent of the CEF (Fig. 1–3). These findings are in agreement with the recently published results by Capó-Bauçà et al. (2022) for submerged angiosperms. Hence, it appears that *C. braunii* primarily relies on passive CO_2_ diffusion for carbon acquisition whereby HCO_3_^-^ uptake plays a rather minor role. In this respect, it is important to consider the distinctive membrane properties of *Chara*,including the pH-based inhomogeneities, containing patterns of alkaline and acid regions along the cell (“pH banding”) and charasomes (Beilby & Bisson, 2012; Foissner et al., 2015). Hereby, the internodal acidification of the surface could lead to the conversion of HCO_3_^-^ to CO_2_ to increase local CO_2_ concentration and diffusion into the cell to increase photosynthetic efficiency (Beilby & Bisson, 2012; Eremin et al., 2019). The identity of proposed mechanisms might vary greatly between different *Chara* spp. (Baastrup-Spohr et al., 2015), and the previously shown efficient HCO_3_^-^ uptake might not fill the whole picture.

Another possibility, which was not elucidated in this paper is the involvement of CAs in the acquisition mechanism of *C. braunii*. The activity of periplasmic CAs to dehydrate HCO_3_^-^ to CO_2_ has previously been shown for *C. tomentosa* (Ray et al., 2003) and hence could also be involved in the carbon uptake mechanism of *C. braunii*. Furthermore, in preliminary long-term incubations in our lab, a repressive effect of NaHCO_3_^-^ on *C. brauniis* growth was observed, enhancing the hypothesis of a minor role of direct uptake of HCO_3_^-^ in this alga.

To gain additional wide-ranging insights into the carbon acquisition mechanisms across the genus *Chara, C. braunii* presents an excellent candidate for further evaluation. This *Chara* appears more broadly related to the genus *Nitella* regarding its ecological niche (Schubert et al., 2018) and its missing cortication distinctively separates it from the rest of the *Chara* genus (Bachy et al. 2022; Raven et al., 2012). (Becker et al., 2016). Hence it poses as an interesting candidate for freshwater algae to aid in further unraveling and improving our understanding of basic pathways as most eukaryotic algae present a CCM, its origin remains less clearly understood, which continuously holds true for *C. braunii*

## Acknowledgments

We thank Dr. Christian Porsche (University of Rostock) for advice on the statistical data treatment.

## Author contributions

HS and MH designed the study, CMH performed and evaluated most experiments, HS evaluated PAM measurements, HS and MH supervised experiments, CMH and MH wrote the manuscript with input from all authors.

